# Geographic Patterns of Bacterioplankton among Lakes of the Middle and Lower Reaches of the Yangtze River Basin, China

**DOI:** 10.1101/536219

**Authors:** Chengrong Bai, Jian Cai, Lei zhou, Xingyu Jiang, Yang Hu, Jiangyu Dai, Keqiang Shao, Xiangming Tang, Xiangdong Yang, Guang Gao

## Abstract

In aquatic ecosystems, microbial biogeography research is critical for unveiling the mechanisms of microbial community succession. However, little is known about the microbial biogeography among interconnected lakes. To address this deficit, we used high-throughput sequencing to explore geographic patterns and the relative importance of ecological processes that shape these patterns in abundant and rare bacterial subcommunities from 25 lakes across the middle and lower reaches of Yangtze River basin (MLYB, located in southeast China), where most of the lakes are interconnected by river networks. We found that there were significant differences in both abundant and rare bacterial subcommunities between the two lake groups that were far from each other, while were no difference among the nearby lakes in each group. Both abundant and rare bacteria followed a strong distance-decay relationship, especially for rare bacteria. These findings suggest that although the interconnectivity between lakes breaks the geographical isolation of bacteria, the dispersal capability of bacterial taxa was still limited by geographic distance. We also found that although deterministic processes and stochastic processes together drive the bacterial subcommunities assembly, the stochastic processes (based on adjusted *R*^2^ in redundancy analysis) exhibited a greater influence on bacterial subcommunities. Our results implied that bacterial dispersal among interconnected lakes was more stochastically.

**Importance:** Unraveling the relative importance of ecological processes regulating microbial community structure is a central goal in microbial ecology. In aquatic ecosystems, microbial communities often occur in spatially structured habitats, where connectivity directly affects dispersal and metacommunity processes. Recent theoretical work suggests that directional dispersal among connected habitats leads to higher variability in local diversity and among-community composition. However, the study of microbial biogeography among natural interconnected habitats is still lacking. The findings of this study revealed interesting phenomena of microbial biogeography among natural interconnected habitats, suggested that the high interconnectivity reduced the spatial heterogeneity of bacteria, and caused the dispersal of bacteria to be more stochastically. This study has provided a deeper understanding of the biogeographic patterns of rare and abundant bacterial taxa and their determined processes among interconnected aquatic habitats.

## Introduction

In aquatic systems, bacteria are the major drivers of nutrient regeneration and energy flow (1, 2), and understanding their spatial diversity and community structure is critical for understanding their relationship with ecosystem functioning (3–5). Compared with large animals and plants, bacteria are more complex in their spatial diversities because of their small size, large numbers, and rich biodiversity (6). A typical phenomenon of most bacterial community compositions (BCCs), is that a few species have numerous individuals, while numerous species have relatively few individuals; the latter is often called the “rare biosphere” (7).

Abundant taxa traditionally considered to be the core groups that performed major ecological functions, and these taxa were therefore intensively studied (8, 9). However, more recent studies have suggested that rare bacterial taxa also play a key role in global biogeochemical cycles and ecosystem functions (7, 10–12). Therefore, the biogeography of both the abundant and rare bacteria needs to be explored rigorously.

The major research focus in microbial biogeography is to reveal the biogeographic patterns of microbial taxa, and to explore the driving processes that shape these patterns (9, 13). Although recent high-throughput sequencing (HTS) studies have provided important insights into microbial biogeography (4), there are still many unresolved problems and disputes (14–16).

Debate continues about whether microorganisms have geographic patterns. Early studies suggested that organisms less than 1 mm long did not have biogeographic patterns for their strong dispersal capabilities (6), and the classic hypothesis was interpreted as ‘everything is everywhere’ (17). However, the recent evidence contradicts this, and shows that BCCs are indeed regularly distributed in time and space (16).

A second debate concerns the processes that shape microbial biogeography. Growing evidence supports that both deterministic (environmental and spatial selection-related) and stochastic (dispersal-related) ecological processes determine the BCCs (13). Deterministic processes mean that the composition and relative abundance of species are determined by certain abiotic and biotic factors (18–20). For example, salinity is the main factor influencing the global diversities and spatial distribution patterns of microorganisms, according to Lozupone and Knight (21) and Caporaso et al. (22). Other studies, in aquatic systems, suggest that deterministic processes such as elevation gradient, salinity, and land use, are the main constraints affecting the high-level heterogeneity of bacterial diversities and biogeographic patterns (9, 23–25). Stochastic processes also can explain the biogeographic patterns of bacteria. For instance, Chen et al. (26) revealed that the planktonic community in four marine ecosystems, around Xiamen island in southeast China, was well explained by stochastic processes. At a large spatial scale, Mo et al. (27) reported that in three subtropical bays of China, stochastic processes has a slightly greater influence on both abundant and rare subcommunities than did environmental selection (deterministic processes).

Abundant and rare bacterial taxa are differentiated from each other by two key significant differences that may influence their geographic patterns: (1) They have different community structures and relative abundances, and may experience different limitations to dispersal. For example, Mo et al. (27) revealed that rare taxa have a stronger response to dispersal limitations than abundant bacterioplankton subcommunities in subtropical bays of China. The different result also reported by Wu et al. (28), who found that abundant taxa had greater dispersal limitations in the surface layer of northwestern Pacific Ocean. (2) Abundant and rare taxa carry out different ecological strategies, may have different ecological niches, and are likely to have different responses to environmental changes. For instance, Liu et al. (13) suggested that among lakes and reservoirs of China, the geographic pattern of rare bacteria is more likely determined by environmental conditions, but abundant taxa are mostly governed by spatial factors. In addition, Liao et al. (9) reported that, among 21 lakes in the Yungui Plateau in China, rare taxa are subject to different environmental drivers than those affecting abundant taxa.

In aquatic ecosystems, microbial communities often occur in spatially structured habitats, where connectivity directly affects dispersal and metacommunity processes (29). The phenomenon of habitat connectivity occurs in open waters (oceans, bays, and lakes) and in riverine ecosystems. The extent to which habitats within open waters and rivers are connected longitudinally depends on their position and spatial dynamics (30–32). Some recent simulated works suggest that dispersal along connected habitats (e.g., dendritic landscapes) often biased the hydrodynamic direction. This biased connectivity leads to a great spatial heterogeneity of species (32, 33).

In natural conditions, however, some habitats are interconnected with each other without significant bias. Few studies have reported the biogeographic patterns of bacteria among interconnected aquatic habitats, and these studies are focused on the ocean or the marine bays (26, 27). The bacterial biogeography among interconnected lakes is still little known. Therefore, understanding the diversity and biogeographic patterns of bacteria among interconnected lakes will aid our understanding of microbial biogeography, and reveal more information about the ecological processes and mechanisms that underlie and maintain ecosystem function in the terrestrial-aquatic ecosystems.

An intriguing area for study is in the east of China, namely the middle and lower reaches of the Yangtze River Basin (MLYB), which is low and flat, and covered by hundreds of shallow lakes and intricate river networks (34). Some of the nearby lakes are connected by river networks, especially in the wet season. More importantly, with the frequent change of regional water level in the wet season, there is a mutual hydraulic exchange among these lakes. Considering the special relationship among the lake habitats, in our present study we hypothesize that: (1) the high connectivity within and among lakes reduces regional differences of geographic patterns of abundant and rare bacterial taxa, but the bacterial dispersal still limited by geographical distance, (2) the “rare biosphere” with numerous taxa but few individuals in each taxon, are more likely to change in the process of dispersal than are the few taxa with abundant bacteria (35); and (3)in connected lakes, stochastic processes exhibited a greater influence on the geographic pattern of bacteria than do deterministic processes.

## Results

### The division of abundant and rare taxa, and multivariate abundance cutoff

In total, 720,326 high-quality sequences were recovered, with a mean of 28,813 ± 4,218 per sample, which clustered into 9,847 OTUs. At regional and local levels, 468,822 sequences (65.1%), representing 81 (0.8%) OTUs were classified as abundant, while 21,083 (2.9%) sequences, representing 6,788 (68.9%) OTUs were classified as rare (Table S1).

Among all lake samples, no single OTU was abundant (> 1%) in all samples; only two OTUs (including 97,781 sequences) with > 1% abundance were present in > 70% of the samples, and 49 OTUs (including 26,319 sequences) were classified as being abundant in only one lake. Among all lake samples, 6,238 OTUs (91.9% of the total) were classified as always rare, and 3,086 OTUs (including 95,443 sequences) with < 0.01% abundance were present > 70% of the samples (Fig. S1).

The result of MultiCoLA shown that little variation in both the Spearman and Procrustes correlations after removal of more than 40% of the rare part of the original data and the truncated data sets. In addition, when the number of rare types increased to 5%, the data structure of the truncated matrices exhibited a little variation (Fig. S2). This result indicated that our definition of abundant (34.5%) and rare (2.9%) bacteria are reasonable.

### Geographic patterns of the bacterioplankton community

Geographic patterns of abundant and rare bacterial communities had significant positive correlation, both dividing two similar clusters (Fig. S3, *P* < 0.01). We found that seven lakes around Lake Taihu (TH) were clustered together (Cluster1), and the remaining lakes were divided into another cluster (Cluster2). Notably, we found that the two clusters were not divided by the middle and the lower reaches (Fig. 2a, S3). In addition, linear regression analysis showed that with increasing geographic distance, the dissimilarities in both abundant and rare communities were increased. Spearman’s correlation analysis gave a correlation coefficient of 0.587 (Mantel test, *P* < 0.01) between geographical distance and the dissimilarity of all bacterial taxa. The Spearman’s correlation coefficients for abundant and rare bacterial taxa were 0.586 (*P* < 0.01) and 0.438 (*P* < 0.01), respectively (Fig. 2b). This phenomenon also suggested that lakes that were closer to each other presented more similar BCCs than lakes that were more distant from each other. Moreover, within the geographic distance, the community dissimilarity coefficient of rare taxa was larger than that of abundant taxa, which indicated that the structure of rare taxa changes more markedly with increasing distance than is the case for abundant taxa (Fig. 2b)

**Fig. 1.**
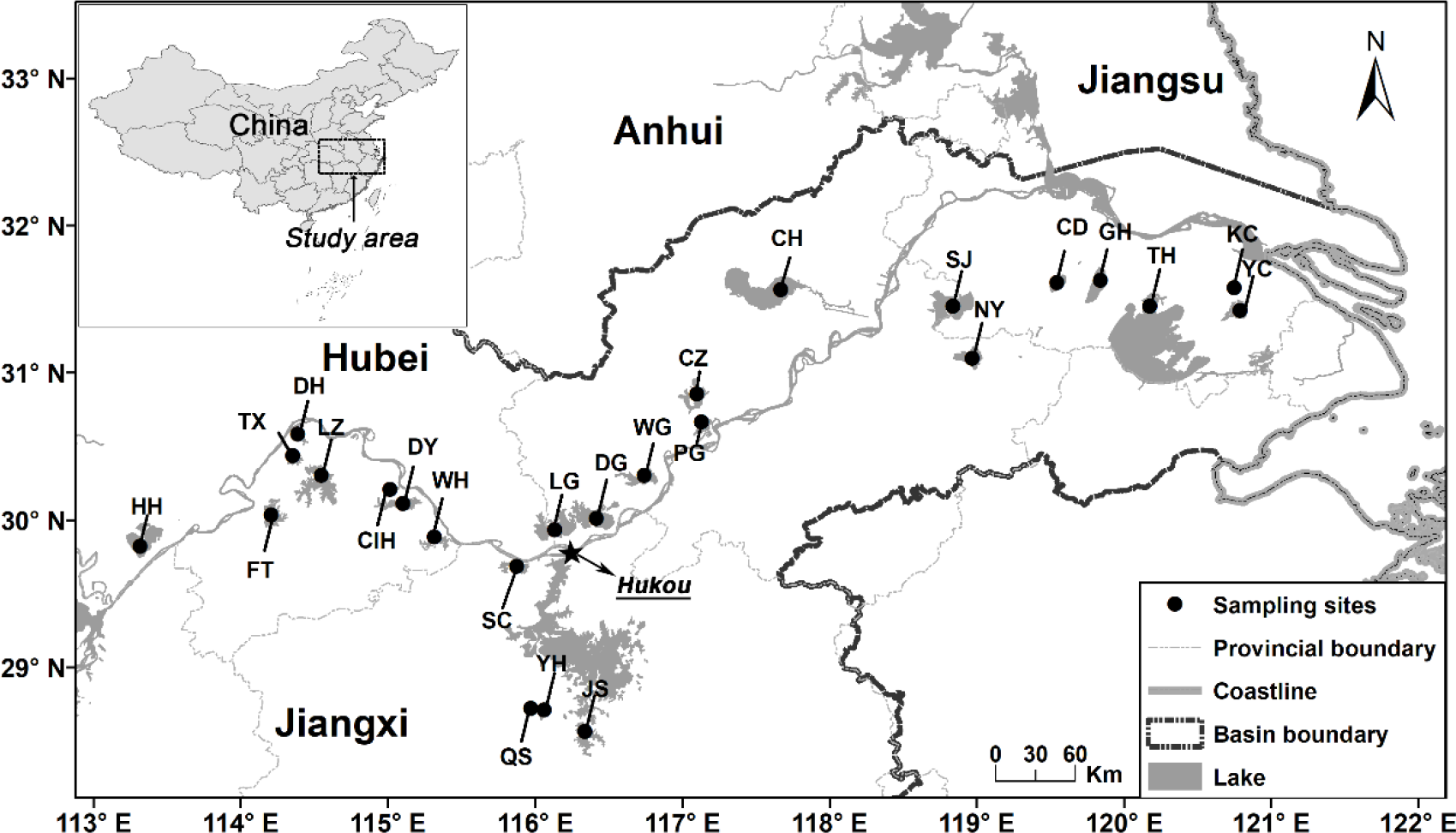
Location of the 25 lakes in the middle and lower reaches of the Yangtze River Basin, China

**Fig. 2.**
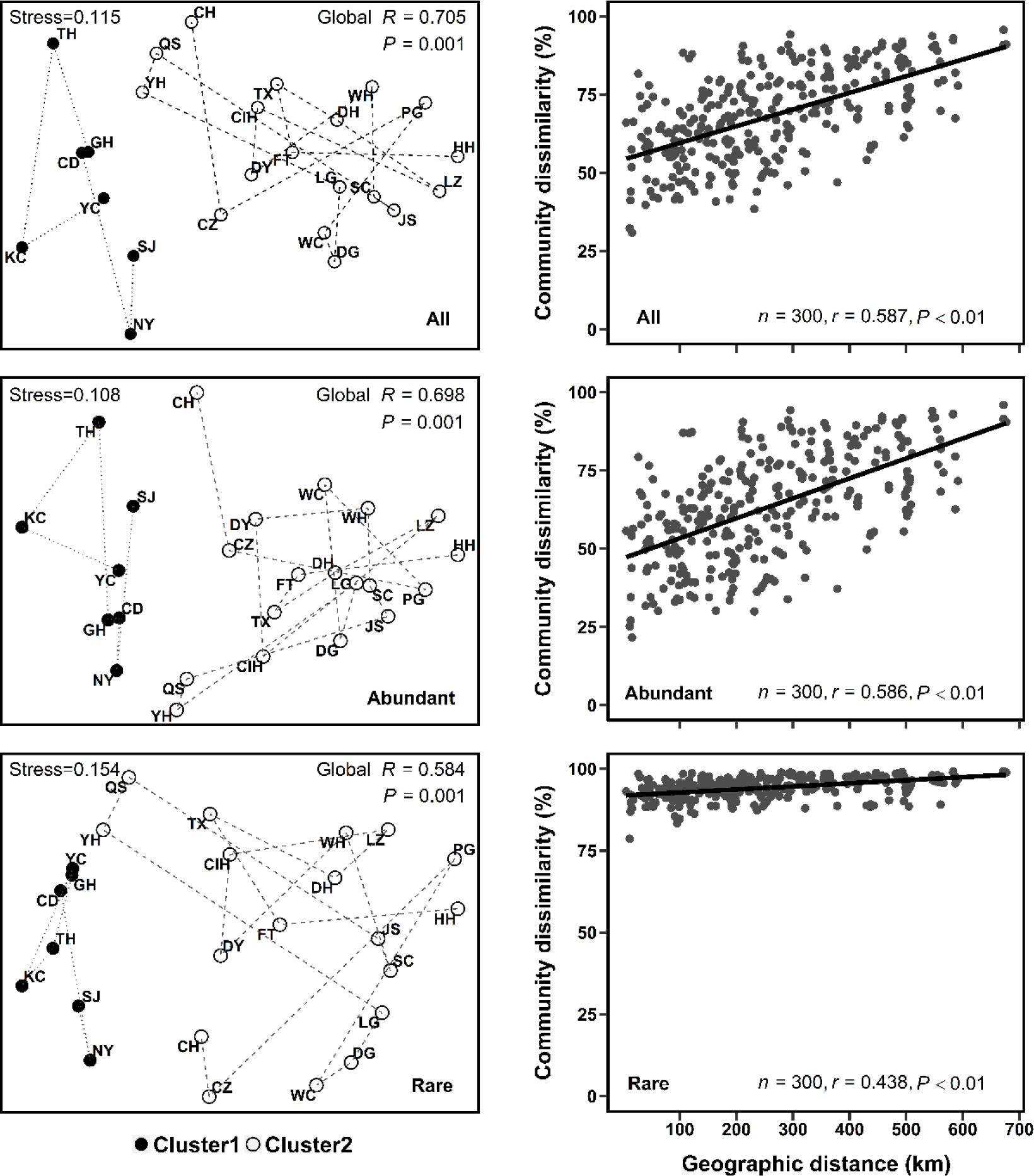
**(a)** The non-metric multi-dimensional scaling (NMDS) for bacterial communities in the 25 lake sampling sites. **(b)** Spearman’s rank correlations between the Bray–Curtis dissimilarity of bacterial communities and geographic distance (*n* is the number of comparisons). All: all bacterial taxa; Abundant: abundant taxa; Rare: rare taxa.

### The relationship between bacterial mean relative abundance and their sites occupied

The bacterial mean relative abundance, for both abundant and rare taxa, was significantly positively correlated with sites occupied (abundant taxa *r* = 0.409, rare taxa *r* = 0.693, *P* < 0.01) (Fig. 3). The majority of abundant OTUs occupied > 50% of sampling sites, but all rare OTUs occupied < 50% of sampling sites (Fig. 3).

**Fig. 3.**
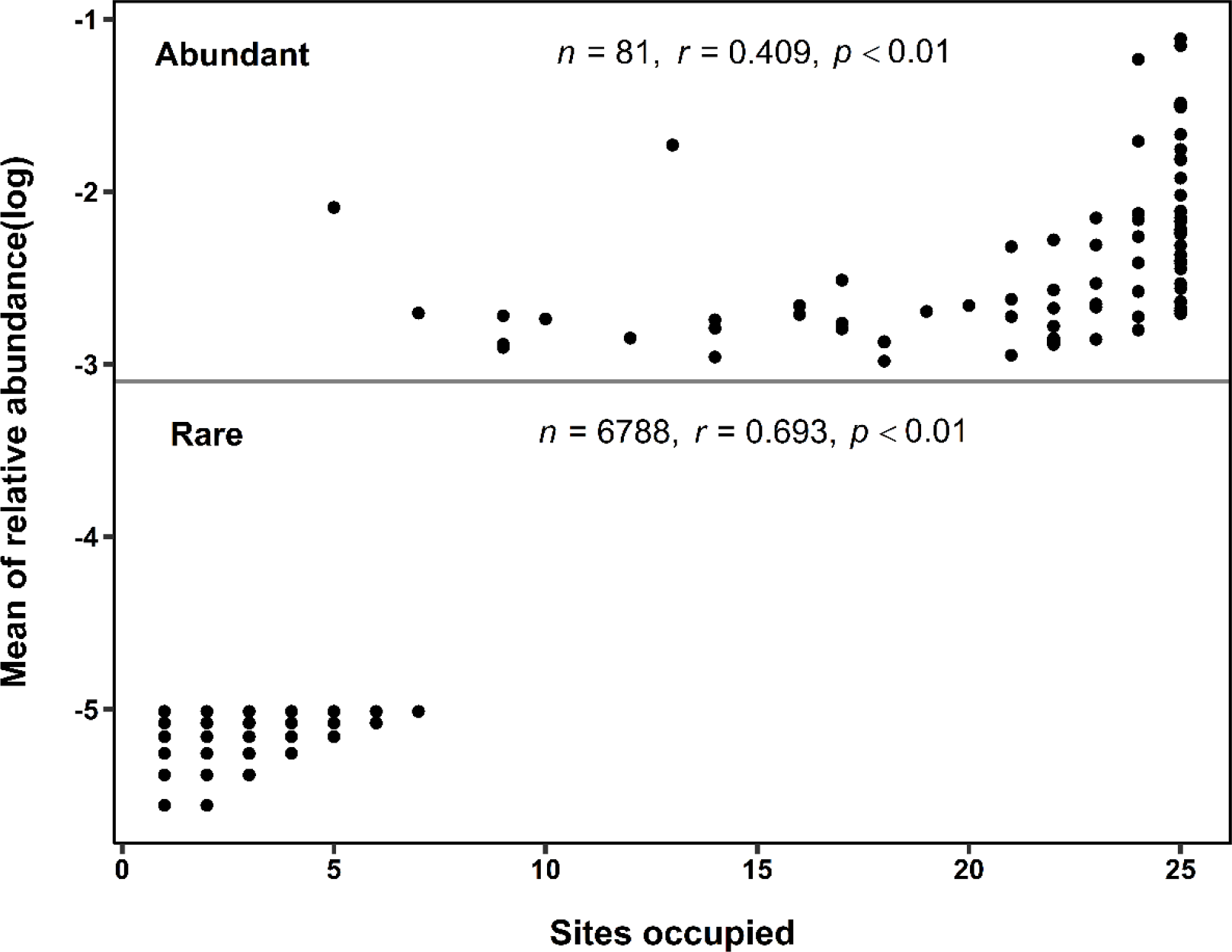
Abundance-occupancy relationship of bacterial taxa. Spearman’s rank correlation between mean relative abundance of abundant bacteria (upper panel) and rare bacteria (lower panel), and number of samples occupied (*n* is the number of OTUs)

### Taxonomic distribution of abundant and rare bacteria

All reads were classified and grouped under 43 phylum-level taxonomic groups (including 2 unknown phyla). Across all samples, *Actinobacteria* and *Proteobacteria* were the two most abundant phyla of bacteria, their average relative abundance accounting for 48.2% and 19.4% of all clean reads, respectively. The next most abundant phyla were: *Verrucomicrobia* (8.1%), *Bacteroidetes* (7.0%), *Planctomycetes* (6.2%), TM7 (4.4%), *Chlorobi* (2.8%), and *Chloroflex* (1.4%). Within the *Proteobacteria* phylum, *α-Proteobacteria* was the most abundant group, accounting for 6.6% of all clean reads; then followed by *β-Proteobacteria*, *γ*-*Proteobacteria*, TA18, and *δ-Proteobacteria*, their relative abundances were 6.4%, 3.5%, 1.5%, and 1.3%, respectively (Fig. S4).

Across all samples, the community structures of both abundant and rare taxa at the phylum level have certain differences. (1) The rare taxa have more species than abundant taxa (Fig. 4). (2) The relative abundances of different phyla in abundant and rare taxa were different. The phyla of the abundant community were dominated by a large group (*Actinobacteria*), while the relative abundance of each rare phylum was relatively uniform. The relative abundance of the same phylum in abundant taxa and rare taxa was as follows: *Actinobacteria* (abundant 60.7 ± 10.4% vs rare 24.9 ± 8.1% (mean ± s.e.)), *Verrucomicrobia* (10.0 ± 7.6% vs 4.2 ± 2.0%), *α-proteobacteria* (3.4 ± 4.4% vs 10.2±3.1%), *β-proteobacteria* (4.4 ± 4.9% vs 9.4 ± 5.3%), *γ-proteobacteria* (2.1± 1.8% vs 6.8 ± 3.1%), *Planctomycetes* (6.4 ± 6.0% vs 8.0 ± 3.7%), *Bacteroidetes* (3.2 ± 3.7% vs 11.1 ± 8.4%), *TM7* (4.5 ± 7.8% vs 2.9 ± 2.5%), *Chlorobi* (3.5 ± 4.0% vs 3.7 ± 2.9%), and *TA18-proteobacteria* (1.3 ± 1.7% vs 2.3 ± 2.2%). Moreover, there were several phyla that showed high relative abundance in rare taxa only, such as *δ-proteobacteria* (5.2 ± 2.9%), *Acidobacteria* (1.9 ± 1.3%), and *Firmicutes* (1.5 ± 0.8%). And a total of 5.4±3.1% groups were occupied by large numbers of low abundance phyla (Fig. 4).

**Fig. 4.**
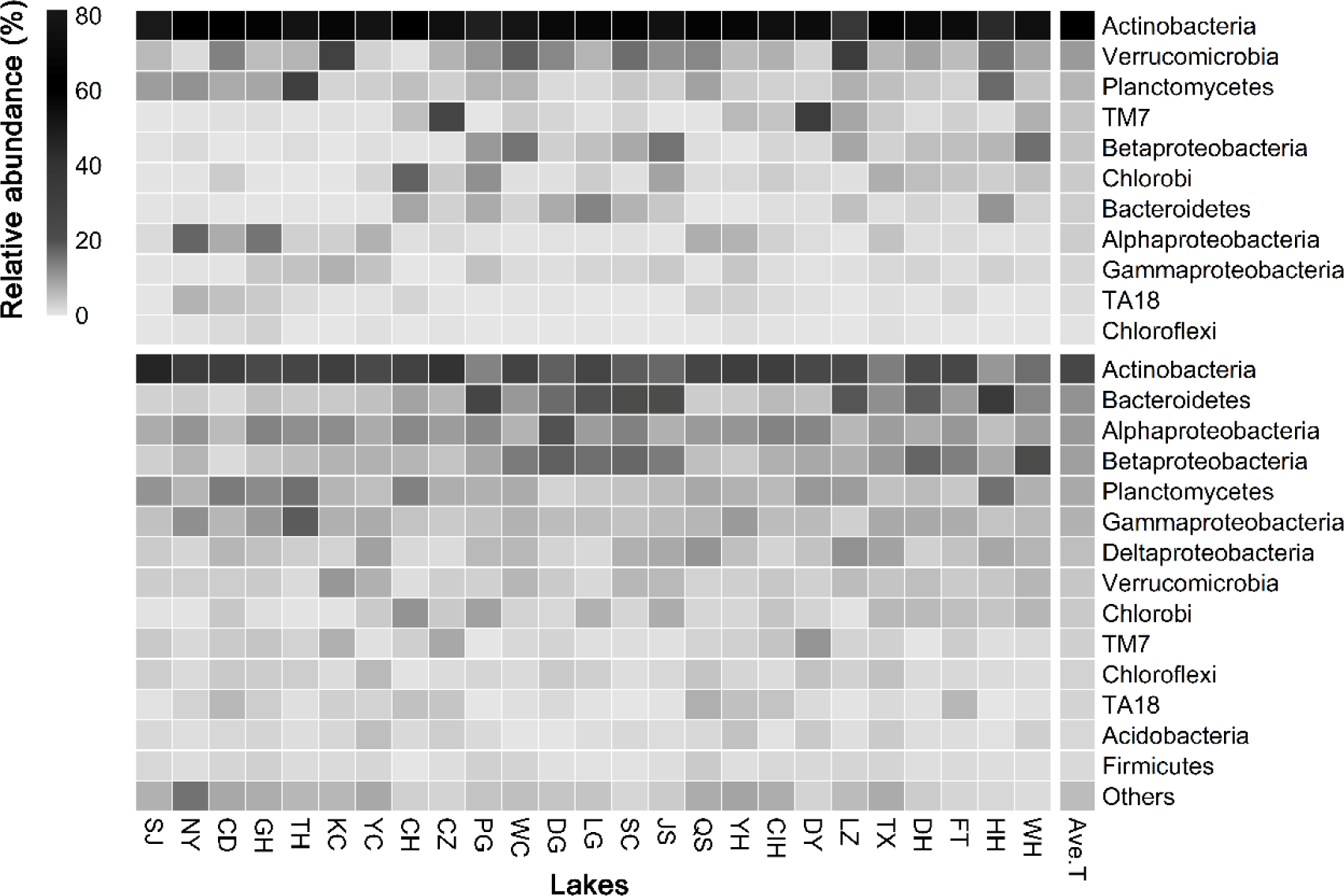
Heat map showing community composition of abundant bacterial taxa compared with rare bacterial taxa in 25 lakes of MLYB, China

### The relative contribution of deterministic and stochastic processes in the assembly of abundant and rare taxa

After analysis, 14 physicochemical variables (Table S2) and 6 significant PCNMs (PCNM1-6, *P* < 0.05) were subject to RDA, CCA, and VPA analyses. The Mantel results revealed that both environmental variables and spatial variables had a significant effect on bacterial sub-communities. Abundant bacterial communities were primarily driven by environmental variables, contrasting with rare bacterial sub-communities which were primarily driven by spatial variables (Table 1).

**Table. 1.** Mantel tests for the correlations between community similarity and local environmental and spatial variables (using Spearman’s coefficient)

| Effects variables | All bacteria | Abundant bacteria | Rare bacteria |
| --- | --- | --- | --- |
| Environmental variables | 0.420 *** | 0.350 *** | 0.349 *** |
| Spatial variables | 0.385 *** | 0.321 *** | 0.464 *** |

To calculate the similarity matrix, the Euclidean distance was used. Significance was tested based on 999 permutations. \*\*\* $P < 0.001$

The RDA ordination showed that the change of the abundant bacterial sub-community was significantly related to two environmental variables (Temp and Chl *a*) and one spatial factor (PCNM1) (by forward model selection) (Fig. 5a). In contrast, the changes in the rare bacterial sub-community taxa were significantly related to the same environmental variables (Temp and Chl *a*) but more spatial factors (PCNM1and PCNM2) (Fig. 5a).

**Fig. 5.**
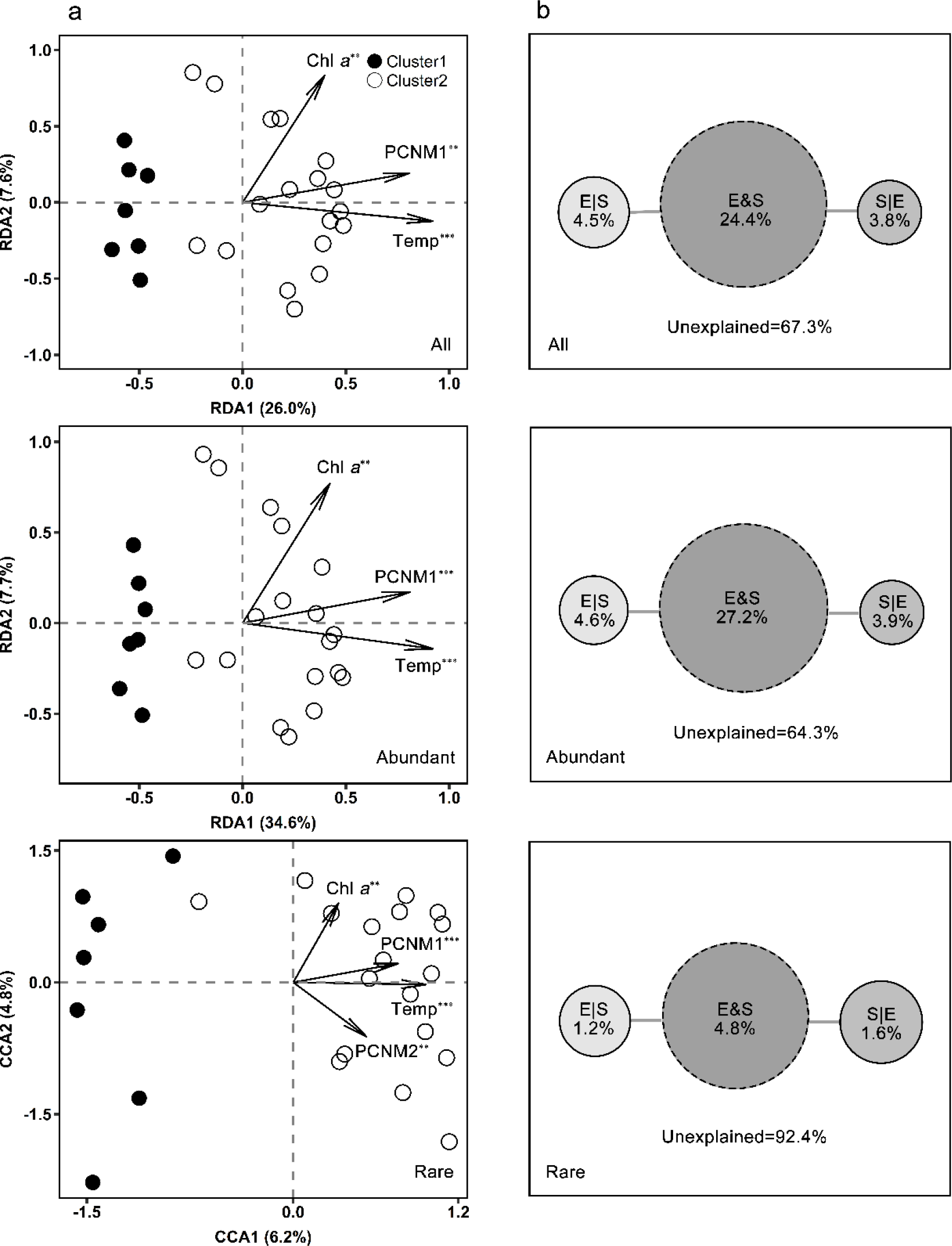
**(a)** Ordination plots showing all, the abundant, and the rare BCCs in relation to significant local environmental variables and spatial variables; significance * *P* < 0.05, ** *P* < 0.01, \*\*\**P* < 0.001. **(b)** Variation in all, the abundant, and the rare bacterioplankton metacommunity explained by environmental and spatial and variables. E|S indicates pure environmental variables, S|E pure spatial variables, E&S share explained variation. Unexplained = 1-E|S-S|E-E&S. All, Abundant, and Rare represent entire taxa, abundant taxa, and rare taxa, respectively.

The result of VPA showed that deterministic processes, namely environmental variables (E) and spatial distance (S), accounted for 35.7% of the abundant bacterial taxa variations; the part accounted for by E and S together was 27.2%. Abundant bacterial taxa were determined by environmental variables, which accounted for 31.8% of the abundant taxa variation (Fig. 5b). For rare bacterial taxa, however, deterministic processes (including E and S) only accounted for 7.2% of the variations. The spatial variable was the dominant driver, explaining 6.4% of the rare taxa variation (Fig. 5b).

In addition, the S alone can explain 3.9% and 1.6% of the abundant and rare variation, respectively. It was notable that a large proportion of bacterial taxa variations were not explained by deterministic processes, especially for rare species.

To explore the effects of the stochastic process on bacterial geographic patterns, the neutral model was employed in our study. The result showed that the neutral model explained a large fraction (*R*^2^ = 52%) of the variation of all bacterial taxa (Fig. 6).

**Fig. 6.**
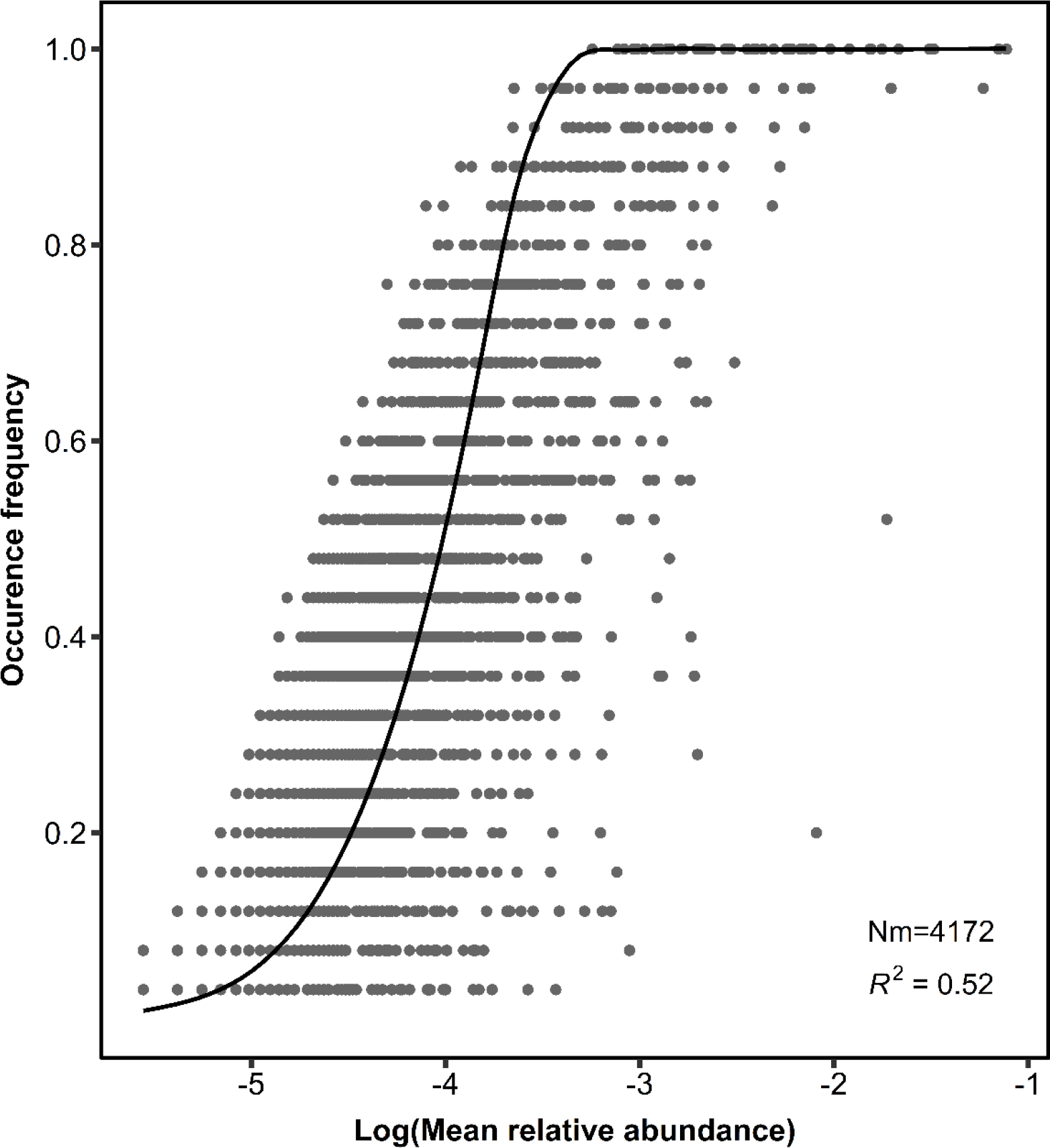
Fit of the neutral community model (NCM) of community assembly. The predicted occurrence frequencies are shown for all bacterial taxa. The solid black lines indicate the best fit to the NCM as in Sloan et al. (2006). *NM* indicates metacommunity size times immigration, *R*^*2*^ indicates the fit to the *NM*

## Discussion

### Geographic patterns in abundant and rare bacteria

The study of BCCs and their biogeographic patterns in particular areas is a fundamental research area in aquatic microbial ecology (1), in which bacterial biogeography considers the relationship between the biodiversity patterns and geographic distributions. New molecular tools and technologies have revealed more information about the “rare biosphere” to us, and rare bacteria are now recognized as having a great effect on ecological functions and mechanisms (12, 36). The ecological relationship between abundant bacteria and rare bacteria, especially the correlation between their geographical patterns has become an attractive research area.

Habitat connectivity directly shapes diversity patterns in microcosm metacommunities at different levels (29). Recent theoretical studies of bacterial biogeography have considered both isolated and interconnected habitats. Among isolated habitats, some studies have reported that the division of bacterial biogeography was consistent with geographical regions. For instance, Liu et al. (13) revealed that in lakes and reservoirs in different regions of China, geographical regions were significantly separated for BCCs. Logares et al. (4) also found that the regional bacterial community was mainly structured by geographic origin. Among interconnected habitats, bacterial subcommunity is complicated among the different regions or sites. For example, Tang et al. (1) reported that free-living (FL) and particle-attached (PA) bacteria were not significantly separated in the different areas of Lake Taihu. At a large spatial scale, namely in some marine bays in China, bacterial biogeography among some interconnected habitats was consistent with their geographical area (26, 27, 37).

Patterns of bacterial abundance are suggested to be the outcome of processes that are inherently driven by the structure of the respective habitat network [30, 31]. Our observation provides important insights for exploring bacterial community patterns among interconnected habitats. Our observations have shown that all lakes were significantly separated as two large clusters at *P* = 0.001 for the bacterioplankton communities. Interestingly, we found that there is a relatively large geographic distance between the lakes in cluster1 and cluster2 (Fig 2a). In this scenario, the dispersal of bacteria between the lakes in cluster1 and cluster2 may be more difficult. A similar phenomenon also reported in some large-distance marine bays in China (27).

However, the ordination of the lakes in each cluster is scattered (Fig 2a). In MLYB, many nearby lakes are connected by intricate river networks, which provide pathways for bacterial dispersal, and break the geographical isolation among lakes. In particular, our samples were collected during the wet season, when high water levels ensured that most lakes were fully connected through the river networks. Furthermore, there was no significant hydrodynamic bias among these lakes, as they are locating in plain areas. In this scenario, the dispersal of bacteria among different lakes is likely to be more randomly. More importantly, contrary to the biased connectivity, the spatial heterogeneity of bacterial diversities among interconnected lakes is reduced (32, 33).

Further, some studies have reported that the rare bacterial biosphere may be significantly correlated with abundant taxa, (rather than being a random collection of taxa) (9, 13, 27). The results of our present study, in particular, the correlation of geographic patterns between abundant and rare bacteria, supported this contention: we found similar biogeographic patterns of rare and abundant taxa (Fig. S3, r=0.873, *P* < 0.01). Our present study indicates that even in interconnected lakes, rare bacteria might in a similar manner as abundant ones in responding to dispersal processes or environmental changes (9, 35).

We also report that in the MLYB, both abundant and rare bacterial taxa had a significant distance-decay relationship (Fig. 2b). Some recent studies reported that the distance-decay relationship of bacteria occurs among both isolated habitats (9, 13, 23) and connected habitats (27). Bacterial community dissimilarity rises with increasing geographic distance if: (1) species filter by local habitat conditions, or (2) if they are limited in dispersal (38). Among interconnected habitats, species can disperse directly through connecting channels. However, the distance-decay relationship showed in both the present study and others (e.g. Mo et al. (27) and Wei et al. (26)), still revealed a phenomenon of limited dispersal. Notably, consistent with Mo et al. (27), we found that there was no significant increase in environmental variability with geographic distance. This result may be explained by bacterial individuals tending to colonize nearby sites (38). In this scenario, the distance-decay relationship would be directly related to the spatial scales among habitats, and would still emerge among different habitats with the same environmental conditions or niche requirements (39). The geographic distance between the lakes in each cluster was less than that between the three marine bays studied by Mo et al. [26]. Correspondingly, the smaller is the geographic distance between regions, the higher is the similarity of the community composition. The small geographic distance between lakes in each cluster may explain why our results are similar to those of Tang et al. [11]. On the contrary, the geographic distance between the lakes in cluster1 and cluster2 is much larger. The large spatial distance might lead to differences of bacterial subcommunities between the two regions.

We also found that the community dissimilarity increased more significant for rare bacteria with the increasing of geographic distance than it did for abundant bacteria. Our result was consistent with Mo et al. (27) and Liao et al. (9) who reported that most of the abundant bacteria appeared in most of the sampling sites, while all rare bacteria appeared in only a few sampling sites (Fig. 3). Further, consistent with Mo et al. (27), we found that the biodiversity patterns between abundant and rare taxa are largely different (at phylum level) (Fig. 4). The abundant taxa showed relative few community compositions and high relative abundance, whereas the rare taxa had complex community composition and low relative abundance (Fig. S1, Fig. 4). Liu et al. (13) suggested that rare bacteria with low local abundance have an increased probability of local extinction, and a decreased probability of dispersal.

### Controlling factors for geographical distributions of abundant and rare bacterial taxa

Investigating the controlling factors, and their contributions to geographical distributions, of abundant and rare bacterial taxa are fundamental research topics for ecologists. Improved understanding of these topics will help us to reveal the reasons for particular biogeographic patterns, and allow prediction of the community succession in the habitat. Study of relevant deterministic processes may focus on ecological selectors as limiting factors, including nutrients, land use, and habitat connectivity (1, 9, 29). However, spatial variation in the BCCs is also influenced by stochastic processes, such as dispersal limitation, successional stages, and random effects (40).

Previous research works have highlighted two key areas for which further study is needed (27, 41). First, few studies have investigated the influence of both deterministic and stochastic processes on the biogeographic pattern of bacteria. Second, the unexplored topic of the influence of both deterministic and stochastic processes on the biogeographic pattern of bacteria in interconnected lakes. So, to address these deficits, in our present study, we focused on these unexplored problems, and tried to reveal the rules of bacterial biogeography among interconnected terrestrial waters.

We used ecological statistical methods (forward selection) to verify the correlation between deterministic variables (environmental and spatial variables) and the spatial distribution of abundant and rare bacterial taxa (42). We found that only two environmental factors (Temp, Chl *a*) were significantly related to the spatial distribution of abundant and rare taxa subcommunities (Fig. 5a). These environmental variables exactly reflected the habitat requirements of bacteria in this area. Most of the lakes in the MLYB are suffer from mesotrophic or eutrophic (43), with extremely high phytoplankton and Chl *a* values. The formation of phytoplankton directly provides energy and materials for the growth and metabolism of bacteria (1), so that determines the bacterial biodiversity. In addition, the temperature is critical for maintaining bacterial growth and metabolic activity. Therefore, the change in temperature systematically altered the relationship between biodiversity and ecosystem functioning (44–46). We notice that although our samples were collected at the same time, the surface water temperature has changed greatly during the rainfall process. Across all lakes, the maximum temperature difference reached 9.55°C (Table S2). Large temperature difference might be another important reason for the bacterial spatial heterogeneity.

We found that both environmental and regional variables were significantly related to community structures of abundant and rare bacterial taxa (Table 1), and that these two variables together explained a large proportion of bacterial distributions (Fig. 5b). The spatial variables explained more variation of rare bacterial community than did the environmental variables (Fig. 5b). In contrast, the environmental variables explained more variation of the abundant bacterial community than did the spatial variables (Fig. 5b). This result was consistent with the Mantel tests for the correlation between community similarity and local environmental and regional factors using Spearman’s coefficient (Table 1).

We note that a contrasting result was reported from marine bays by Mo et al. (27). These authors reported that while environmental and spatial variables together explained a large part of the variations in abundant communities, they explained little of the variations in rare communities. They suggested that purely environmental components were slightly less influential contributors than the purely spatial variables in regulating the assembly of both abundant and rare subcommunities.

The discrepancy between our present work and that of Mo et al. [26] may be explained in two aspects. Firstly, we hold that interconnectivity among lakes leads to a frequent exchange of energy and environmental variables, and reduces the heterogeneity of environment among nearby lakes (Fig. S5). Secondly, with low relative abundance, the rare taxa, were influenced by spatial variables more strongly than were the abundant taxa (Fig. 2b).

In addition, we hold that the interconnectivity among lakes leads to the dispersal of bacteria in different habitats to be more stochastic than deterministic, especially for rare bacteria. Thus, the contribution of environmental and spatial variables to the geographic pattern of bacterial communities were quite low, especially for rare species (Fig. 5b). Our result was consistent with several recent studies on different interconnected marine bays, and their abundant and rare subcommunities (26, 27, 37). However, in contrast to the isolated lakes, which indicated that abundant and rare bacterial taxa were strongly influenced by the deterministic process, and only weakly influenced by the stochastic process (9).

The NCM analysis showed that stochastic process explained a large fraction (*R*^*2*^=52%) of the variation of all bacterial taxa in the interconnected lakes in the MLYB (Fig. 6), consistent with similar studies on the bacterial subcommunities of interconnected marine bays (26, 27, 37), but contrasting with the situation in isolated lakes (9). These results indicate that the interconnectivity among habitats increases the randomness of bacterial dispersal. Two main reasons could support our result: (1) most of the lakes were interconnected by numerous river networks so that the flow of water among these lakes was not a significant bias; and (2) bacteria cannot avoid adverse conditions or seek favorable environments, in contrast to larger aquatic animals. Moreover, this result was also consistent with our finding of the no regional differences of bacteria. In addition, there was still a large amount of unexplained variation that could be attributed to other biotic or abiotic variables than the ones mentioned in this paper.

## Conclusions

This study has provided a deeper understanding of the biogeographic patterns of rare and abundant bacterial taxa and their determined processes among interconnected aquatic habitats. Our conclusions strongly support our hypotheses, we interestingly found that the high connectivity within and among lakes reduces regional differences of geographic patterns of abundant and rare bacterial taxa, especially among nearby lakes. However, there were significant differences in both abundant and rare bacterial subcommunities between the two lake groups that were far from each other. This phenomenon was consistent with the result that both abundant and rare bacteria followed a strong distance-decay relationship. In addition, the “rare biosphere” with numerous taxa but few individuals in each taxon, were encountered more dispersal limitation and followed a stronger distance-decay relationship. Furthermore, deterministic processes and stochastic processes together drive the bacterial subcommunities assembly, but stochastic processes exhibited a greater influence on the biogeography of bacterial communities among the interconnected aquatic ecosystem.

## Materials and methods

### Study area and sample collection

With a total area of 18,400 km^2^, the MLYB has more than 600 lakes larger than 1 km^2^ (47). These lakes are vital to both regional economic development and ecological stability (48, 49). However, due to heavy external pollutant inputs from local and upstream areas, most of these lakes are polluted, and many are eutrophic (50). There is a strong hydraulic disturbance and exchange among lakes, due to the area’s location in the subtropical climatic zone, with distinct dry and wet seasons.

The MLRB is divided into the middle reaches (MR) and the lower reaches (LR) by the city Hukou (51). Field work was conducted in July 2016. We randomly selected 25 lakes in the MR and the LR from which to collect water samples (Fig. 1, and Table S1). At each lake, one surface water sample (from the top 50cm of the water column) was collected with a 5 L sterilized polypropylene bottles at the lake center. All water samples were transported to the laboratory in dark cooling boxes and processed for physicochemical analyses and bacterial community analyses as soon as possible.

### Physicochemical analysis

Water column depth (WD) and Secchi depth (SD) were measured by a water depth gauge (Uwitec, Austria) and Secchi disk, respectively. Water temperature (Temp), pH, electrical conductivity (EC), dissolved oxygen (DO), salinity (Sal), and turbidity of the epilimnion layer were measured *in situ* with a multiparameter water quality sonde (YSI 6600 V2, Yellow Springs Instruments, U.S.A.). Nine physicochemical parameters were determined using standard methods [40]: total nitrogen (TN), ammonium (NH_4_^+^), nitrate (NO_3_^−^), nitrite (NO_2_^−^); total phosphorus (TP), orthophosphate (PO_4_^3−^); dissolved inorganic carbon (DIC), dissolved organic carbon (DOC), total suspended solids (TSS), and chlorophyll a (Chl *a*).

### DNA extraction, PCR, and high-throughput sequencing

For bacterioplankton analyses, a 100-500 ml water sample was filtered through a 0.22-μm pore size polycarbonate filter (47 mm diameter, Millipore, Billerica, MA, USA) (52), and then the filter was put into a sterile centrifuge tube and rapidly stored at −80℃. Total DNA was extracted directly from the filters, using FastDNA^®^ spin kit for soil (MP Biomedicals, Solon, OH, USA) according to the manufacturer’s instructions. After checked the quality and quantity of the amplified DNA fragment (used the NanoDrop ND-1000 UV/ Vis spectral photometer), the V4 region of the bacterial 16 ribosomal RNA gene was amplified by primers 515F (5ʹ-GTGCCAGCM-GCCGCGGTAA-3ʹ) and 806R (5ʹ-GGACTACHVGGGTWTCTAAT-3ʹ) (53) with 20μL of DNA. The PCR amplifications for each DNA sample followed these steps: initial denaturation at 94 °C for 5 min, followed by 25 cycles of 30 s at 94 °C, 30 s at 50 °C, and 30 s at 72 °C; the final step was an extension at 72 °C for 7 min. Subsequently, the total nucleic acids were sent to BGI Co., Ltd (Wuhan, China) for high-throughput sequencing on an Illumina MiSeq instrument (San Diego, CA, USA).

### Bioinformatics analysis

Raw sequence data were processed using the Quantitative Insights into Microbial Ecology (QIIME V. 1.9.1) pipeline (54). After estimating the quality of raw reads by using FastQC (v0.11.5), the Illumina pair-end reads were assembled using FLASH, with a minimum overlap of 10 bases (55). Then, 16S reads were de-multiplexed and quality-filtered following the pipeline and previous research results (56). Next, we checked and then removed potential chimera sequences with a reference-base method using VSEARCH v1.11.1 (57), which implements the UCHIME algorithm (58).

Bacterial phylotypes were identified and assigned to operational taxonomic units (OTUs, 97% cutoff), based on the 13_8 release of the Greengenes database [48]. After taxonomy classification, representative sequences were aligned and further refined using PyNAST (59, 60), and phylogenetic trees were built using FastTree (61). We removed all archaea, chloroplasts, mitochondria, unassigned sequences, and spurious singletons from the OTU table. Finally, to minimize the sequencing depth of different samples, the high-quality OTU table was randomly resampled based on the sample with the smallest sequencing (18,753 reads) using the QIIME pipeline (54).

### Definition of abundant taxa and rare taxa

We defined abundant and rare OTUs based on their relative abundance, considering two thresholds: on the regional and local scale. At a regional scale, OTUs with a mean relative abundance of > 0.1% across all samples were defined as abundant taxa, and those with a mean relative abundance of < 0.001% were defined as rare taxa (4, 13). At the local scale, the thresholds were: abundant if > 1% within a sample, and rare if < 0.01% within a sample (7). To reduce the effect of arbitrariness, we used the multivariate cutoff level analysis (MultiCoLA V. 1.4) as described by Gobet (62).

### Data analyses

The *β-*diversity indices were analyzed by QIIME (V. 1.9.1) as part of the OTUs’ downstream analysis (54). The geographic patterns of both abundant and rare taxa among the lakes were revealed by non-metric multi-dimensional scaling ordination analysis (NMDS). In addition, the cluster analysis was used to test whether rare bacterial taxa share similar biogeography with abundant ones. For NMDS and Cluster analysis, OTU data were initially subject to Hellinger’s transformation (63). The relationships between bacterial community dissimilarity and geographic distance of these lakes were calculated by Spearman’s rank correlations. The dissimilarity of the bacterial community was determined by Bray–Curtis dissimilarity matrix, and the geographic distance was determined by geographic coordinates of each lake. To determine if there was a significant difference in bacterial communities with increasing geographic distance, the Mantel test was used.

To explore the influence of deterministic processes on BCCs, the canonical ordination analyses were used in the R environment using the Vegan package. The deterministic processes including environmental and spatial variables (geographic distance). The normal distribution of the environmental variables was checked using the Shapiro–Wilk test by SPSS (version 22), and these factors were log(*x*+1) transformed (with the exception of pH) to improve normality and homoscedasticity for multivariate statistical analyses. To avoid collinearity, environmental factors with variance inflation factors (VIF > 20) were deleted. The spatial variables were generated by principal coordinates of neighborhood matrices (PCNM) analysis (64), based on the geographical coordinates of each sample.

The correlations between bacterial communities and environmental and spatial variables were analyzed by suitable ordination analysis method determined by the detrended correspondence analysis (DCA). The result of DCA revealed that the longest gradient lengths for both all and abundant bacteria were < 3, indicating redundancy analysis (RDA) is suitable for both communities; while the longest gradient lengths for rare bacteria was > 4, indicating canonical correspondence analysis (CCA) is suitable for rare bacterial community (65).

Environmental variables and spatial variables that significantly explained parts of the variation in BCCs were selected by forward selection. Based on a Monte Carlo test with 999 permutations, only variables that explained a significant (p < 0.05) additional proportion of total variance were included in the subset of forward selected variables (42). In addition, the contributions of environmental variables and spatial variables to geographic patterns of BCCs were assessed by Variation partitioning analysis (VPA).

To evaluate the potential importance of stochastic processes for BCCs, we used the neutral community model (NCM) (66). This model predicts the relationship between the frequency of detecting an OTU and its relative abundance along taxonomic ranks (37). In the NCM, the parameter *Nm* determines the correlation between occurrence frequency and regional relative abundance, with *N* describing the metacommunity size, and *m* being the immigration rate. The parameter *R*^*2*^ determines the overall fit to the neutral model (66). In our present study, all computations for this model were performed in R (version 3.4.4) (R Core Team, 2015) (67).

### Nucleotide sequence accession number

The 16S gene sequences generated in the present study were submitted to the National Center for Biotechnology Information (NCBI) database (http://www.ncbi.nlm.nih.gov) under the accession number SRP173893.

## Acknowledgments

Our present study was supported by the Major Science and Technology Program for Water Pollution Control and Treatment (2017ZX07203-004), the National Natural Science Foundation of China (41621002, 41790423, 41530753, 41571462), the “One-Three-Five” Strategic Planning of NIGLAS (NIGLAS2017GH05) and the Key Research Program of Frontier Sciences, CAS (QYZDJ-SSW-DQC008).

